# A conserved sequence in *Drosophila* Argonaute1 mRNA contributes to stress response via inducing miR-999 degradation

**DOI:** 10.1101/2022.11.16.515324

**Authors:** Peike Sheng, Lu Li, Tianqi Li, Yuzhi Wang, Nicholas M. Hiers, Jennifer S. Mejia, Jossie S. Sanchez, Lei Zhou, Mingyi Xie

## Abstract

MicroRNAs (miRNAs) load onto Argonaute (AGO) proteins and target messenger RNAs (mRNA) to directly suppress gene expression at the post-transcriptional level. However, miRNA degradation can be triggered when there is extended base-pairing between miRNAs and the target RNAs. Such base-pairing can induce confirmational change of AGO and recruitment of ZSWIM8 ubiquitin ligase to mark AGO for proteasomal degradation, while the miRNAs are subsequently degraded. This target RNA-directed miRNA degradation (TDMD) mechanism appears to be an evolutionary conserved mechanism, but recent studies have focused on identifying endogenous TDMD events in the mammalian systems. Here, we performed AGO1-CLASH (crosslinking and sequencing of miRNA-mRNA hybrids) in *Drosophila* S2 cells, with *Dora* (ortholog of vertebrate ZSWIM8) knockout (KO) mediated by CRISPR-Cas9 to identify five TDMD triggers (sequences that can induce miRNA degradation). Interestingly, one trigger residing in the 3′ UTR of *AGO1* mRNA induces miR-999 degradation. CRISPR-Cas9 KO of the *AGO1* trigger in S2 cells and in *Drosophila* elevates miR-999 abundance, with concurrent repression of the miR-999 targets. *AGO1* trigger-KO flies respond poorly to hydrogen peroxide-induced stress, demonstrating the physiological importance of a single TDMD event.

## Introduction

miRNAs are ∼22-nucleotide (nt) noncoding RNA that act as regulators at the post-transcriptional level and play a key role in essential cellular processes. Typically, mature miRNAs associate with AGO proteins and bind to the target mRNAs through the seed region (nucleotides 2-8 from the miRNA 5′ terminus) to degrade and/or inhibit the translation of these RNA targets^1^. The regulation of miRNA biogenesis has been extensively studied, but much less is known about the degradation mechanisms of miRNA. Generally, miRNAs maintain slow turnover, but some miRNAs can be rapidly degraded, with a half-life from less 2 hours to 10 hours^2, 3^. Recent studies have found that extended complementary pairing between miRNAs and target RNAs can induce accelerated degradation of miRNAs, through a mechanism known as target RNA-directed miRNA degradation (TDMD)^4, 5, 6, 7, 8^.

In canonical miRNA–target interactions, miRNAs mostly bind to targets via base-pairing of the seed region, while the miRNA 3′ end is buried in the PAZ domain of AGO and protected from enzymatic attack^9^. However, the miRNA 3′ end is exposed and enables tailing and trimming when the miRNA interacts with a TDMD-inducing target with extended 3′ complementarity^4, 10^. Extensive TDMD base-pairing also promotes broad structural rearrangements of AGO^11^. ZSWIM8 Cullin-RING E3 ubiquitin ligase recognizes this TDMD-associated conformation and interacts with AGO, resulting in ubiquitylation and subsequent proteasomal decay of the AGO to release the miRNA for degradation by ribonucleases^6, 7^. Interestingly, ZSWIM8 promotes TDMD following a tailing and trimming-independent manner^6, 12^.

Multiple targets that can trigger TDMD have been identified, including synthetic targets, viral transcripts, and endogenous protein-coding and noncoding RNAs^4, 5, 13, 14, 15, 16^. The TDMD phenomenon was first discovered by the Steitz laboratory, who found that the viral noncoding RNA HSUR1 (Herpesvirus saimiri U-rich RNAs) can induce degradation of host cell miR-27^5^. At the same time, The Zamore laboratory using synthetic targets RNA found that extended base-pairing of miRNAs with targets at the 3′ end induces tailing and trimming of miRNAs, thus affecting miRNA stability in *Drosophila* and human cells^4^. Subsequent studies identified multiple endogenous TDMD triggers. These include the long non-coding RNA *CYRANO* which can induce the degradation of miR-7, and the 3′ UTR region of *NREP, Serpine* and *BCL2L11* mRNAs which promote the degradation of miR-29b, miR-30b/c and miR-221/222, respectively^13, 14, 15, 16^. In addition, many potential TDMD triggers were identified by high-throughput screening and validated in reporter transfection assays^13, 15, 17^. With the identification of the endogenous TDMD triggers, the characteristics of the trigger sequences were summarized: 1, base-pairing in the 5′ and 3′ end of miRNA with central mismatches; 2, TDMD trigger sequences are highly conserved in different species; 3, The flanking sequence of the TDMD trigger is also required.

In *Drosophila melanogaster*, two different types of small RNAs, small interfering RNA (siRNA) and miRNA, associate with AGOs to regulate gene expression. Typically, siRNAs bind to AGO2 and directly cleave target RNAs with extensive complementarity to siRNA, while miRNAs usually interact with AGO1 to reduce translation and stability of partially complementary target RNAs^4, 18, 19, 20, 21, 22^. In addition, siRNAs loaded with AGO2 are modified to 2′-O-methylation by the methyltransferase Hen1 at the 3’ end, thus protecting siRNAs from trailing and trimming. However, miRNAs bound to AGO1 do not have this modification^23, 24, 25^. Interestingly, even in the absence of 2′-O-methylation at the 3′ end, the siRNAs that have extensive base-pairing with the target RNAs in AGO2 are spared from TDMD degradation^12^.

Many *Dora* (ortholog of vertebrate ZSWIM8)-sensitive miRNAs have been identified, suggesting that these miRNAs are under TDMD regulation^7^. We have previously adopted AGO-CLASH to identify endogenous triggers from multiple mammalian cell lines^13^. To further investigate the regulatory mechanism of TDMD in *Drosophila*, here we performed a global screening of the TDMD trigger in S2 cells by AGO1-CLASH.

## Results

### TDMD trigger identification in *Drosophila* S2 cells by AGO1-CLASH

We performed AGO1-CLASH in *Drosophila* S2 cells, with *Dora* knockout mediated by CRISPR-Cas9 (Figure 1A)^7^. We first confirmed the abundance of *Dora*-sensitive miRNAs in S2 cells including WT, control-KO (scramble), and *Dora*-KO cells. Northern blot results showed that these miRNAs’ abundance was significantly increased in *Dora*-KO cells (Figure 1B). Meanwhile, we analyzed the abundance of miRNA reads between *Dora*-KO and control-KO in AGO1-CLASH libraries. The abundance of guide strand of the *Dora*-sensitive miRNAs significantly increased, but passenger strand remained largely unchanged, supporting the notion that TDMD is specific to mature miRNAs (Figure 1C). With minor modifications compared to our study in mammalian cells^13^, we analyzed the obtained AGO1-CLASH data, focusing on the pairs of triggers matching the TDMD base pattern (described in detail in Materials and Methods) (Figures 1B and 1C)^7^. Subsequently, we screened for high-confidence TDMD hybrids based on three additional criteria: 1. TDMD-like extensive base-pairing pattern between the miRNA and the target RNA; 2. abundance of the miRNA/target RNA hybrid reads in AGO1-CLASH from *Dora*-KO is higher than 100 reads per million (RPM); 3. there is more than 4-fold increase of miRNA/target RNA hybrid reads in AGO1-CLASH from *Dora*-KO compared with control-KO cells.

**Figure 1.**
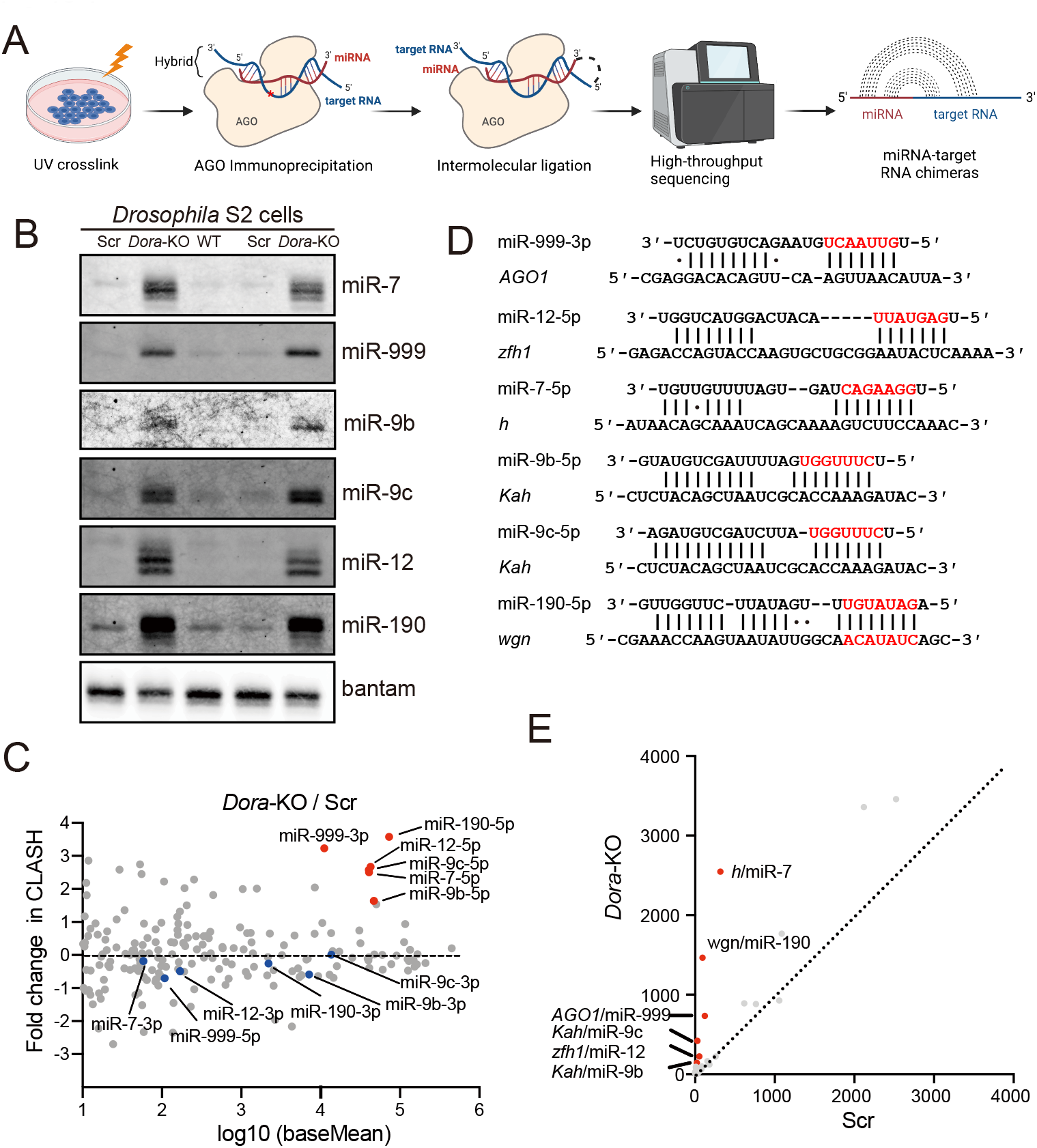
Identification of TDMD triggers in S2 cells by AGO1-CLASH. (**A**) Schematic of AGO1-CLASH. Endogenous miRNAs and target RNAs are crosslinked (254 nm UV) with AGO1 and immunoprecipitated using AGO1 antibody. After AGO1-IP, the miRNAs are ligated directly to their targets to form hybrid molecules for high-throughput sequencing. (**B**) Northern blot analyses of miR-7, miR-999, miR-9b, miR-9c, miR-12, miR-190 and bantam in WT, control-KO (Scramble) and *Dora*-KO S2 cells. The levels of bantam serve as loading control. Changes in miRNA abundance observed from CLASH between *Dora*-KO and control-KO S2 cell. Guide strands of the *Dora*-sensitive miRNAs are indicated by red dots, and the blue dots represent their passenger strands. (**D**) Base-pairing pattern of miRNAs and potential TDMD triggers. Red letters represent miRNA seed region. (**E**) The comparison of potential TDMD miRNA-target RNA hybrids in AGO1-CLASH data obtained from *Dora*-KO and control-KO (Scramble) S2 cells.

In total, we identified five high-confidence TDMD triggers corresponding to *Dora*-sensitive miRNAs in AGO1-CLASH datasets, including *AGO1*/miR-999-3p, *zfh1*/miR-12-5p, *h*/miR-7-5p, *Kah*/miR-9b-5p, *Kah*/miR-9c-5p and *wgn*/miR-190-5p (Figures 1D and 1E, Table S1). Of note, during the preparation of our manuscript, the Bartel group reported the identification of TDMD triggers via prediction through TDMD-like base-pairing patterns and revealed the same set of five triggers from S2 cells^26^. Interestingly, these TDMD triggers all locate in the 3′ untranslated region (UTR) of the mRNAs (Table S1, Figure 2A), the same location as the previously validated endogenous mammalian triggers in mRNAs. It is worth noting that the TDMD trigger of miR-999 is the most conserved among the five triggers. In addition, there are only two nucleotides within the *AGO1* trigger sequence that are not 100% conserved comparing six fly species (Fig. 2A), but neither of them affects the base-pairing with miR-999 (Fig. 1D).

**Figure 2.**
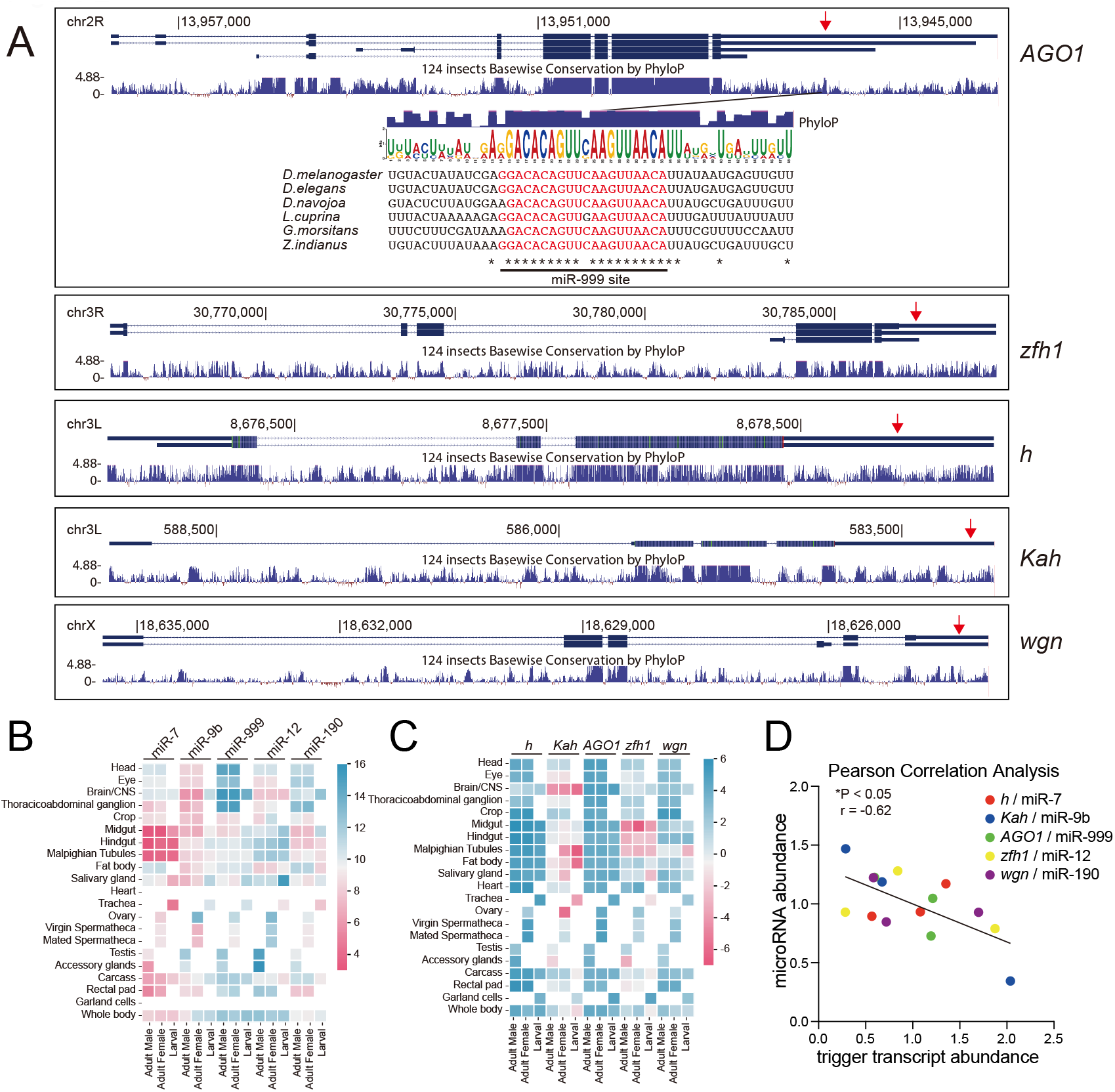
Evaluation of the high-confidence TDMD trigger. (**A**) Genome browser view of trigger transcript models (blue boxes, exons; blue line, introns) with alternative isoforms and conservation plots, which are based on a 124 insects Basewise Conservation by PhyloP (phyloP124way). The transcripts are shown in the 5′ to 3′ direction. Red arrows point to the TDMD triggers. Diagramed below *AGO1* transcript is the trigger site against miR-999 (in red), flanked by 15 nt on each side. The sequence logo based on 6 homologous sequences is shown from the indicated species. Asterisks indicate bases conserved in all of the representative examples. Tissue-specific miRNA abundance (**B**) and trigger transcripts abundance (**C**) in *Drosophila* from FlyAtlas 2. The gene abundance scale (log2 FPKM for genes, log2 RPM for miRNA) is shown to the right of the heatmap. (**D**) Negative correlation between the miRNA abundance and trigger transcript abundance in *Drosophila* whole body. The linear regression is shown with solid line. r, pearson correlation coefficient. *p < 0.05 by two-tailed test.

To gain further support of the TDMD pairs, we examined whether the levels of the five miRNAs and their corresponding TDMD trigger transcripts have negative correlation in *Drosophila*. With the miRNAs and trigger transcript abundance information in different organs of the flies obtained from FlyAtlas 2 database (flyatlas2.org), we observed many instances of reverse correlation between the miRNAs and the trigger transcripts (Figures 2B and 2C). We further normalized triggers and miRNA abundance in the whole body with the average abundance across different organs, a negative correlation is observed based on the Pearson correlation analysis (Figure 2D). These data further suggest the high confidence of our candidate TDMD triggers.

We next examined miRNA abundance in flies with *Dora* mutation to check whether the *Dora*-sensitive miRNAs in S2 cells are also under TDMD control in the animals. Two lines with *Dora* mutation were obtained at the Bloomington *Drosophila* Stock Center [stock number 52333 (y[1] w[*] *Dora*[A] P{ry[+t7.2]=neo FRT}19A/FM7c, P{w[+mC]=GAL4-Kr.C}DC1, P{w[+mC]=UAS-GFP.S65T}DC5, sn[+]) and 52334 (y[1] w[*] *Dora*[B] P{ry[+t7.2]=neoFRT}19A/FM7c, P{w[+mC]=GAL4-Kr.C}DC1, P{w[+mC]=UAS-GFP.S65T}DC5, sn[+])]. Since the homozygous *Dora* mutations are lethal, both lines are heterozygous. Utilizing the green fluorescent protein (GFP) marker on the balancer chromosome, we isolated homozygous *Dora* mutants and control embryos at 10-16 hours after egg laying. We extracted total RNA from homozygous *Dora* mutant and control embryos and performed small RNA sequencing (small RNA-seq). Interestingly, the *Dora*-sensitive miRNAs in embryos are completely different compared with S2 cells, with the miR-310 and miR-3 family members being elevated the most in *Dora* mutant embryos (Figures S1). Such stark difference highlights the potential importance of TDMD regulation in *Drosophila* embryogenesis. The recent paper from the Bartel group has demonstrated that TDMD events targeting miR-310 family members in the *Drosophila* embryo are required for proper embryonic development^26^ . In this study, we followed up the TDMD triggers identified from *Drosophila* S2 cell.

### Validation of TDMD trigger in S2 cells with CRISPR-Cas9 knockout

We next wanted to validate candidate TDMD triggers can affect the abundance of their corresponding miRNAs. To this end, we designed two sgRNAs targeting both sides of the TDMD trigger and cloned the sgRNAs into the pAc-sgRNA-Cas9 plasmid^27^. Pairs of two plasmids targeting each trigger were co-transfected in S2 cells. For each trigger KO, two pairs of sgRNAs were used to obtain two different cell lines (Table S2). Stable polyclonal cells were obtained after selection for one month with 5 µg/mL puromycin. From these stable cells, we extracted total RNAs and performed northern blot analyses for miRNAs. After knocking out the TDMD trigger of *AGO1, zfh1, h, Kah* and *wgn*, the abundance of the corresponding miRNAs all significantly increased, except for miRNA-9c (Figures 3A and S2). Interestingly, compared with other miRNAs, when the TDMD trigger of *AGO1* was deleted, the abundance of miRNA-999 increased to a level similar to that of *Dora*-KO, suggesting that *AGO1* may be the only trigger for miR-999 in S2 cells. Because our TDMD trigger KO cells are polyclonal, we cannot assess whether other sensitive miRNAs have additional triggers.

**Figure 3.**
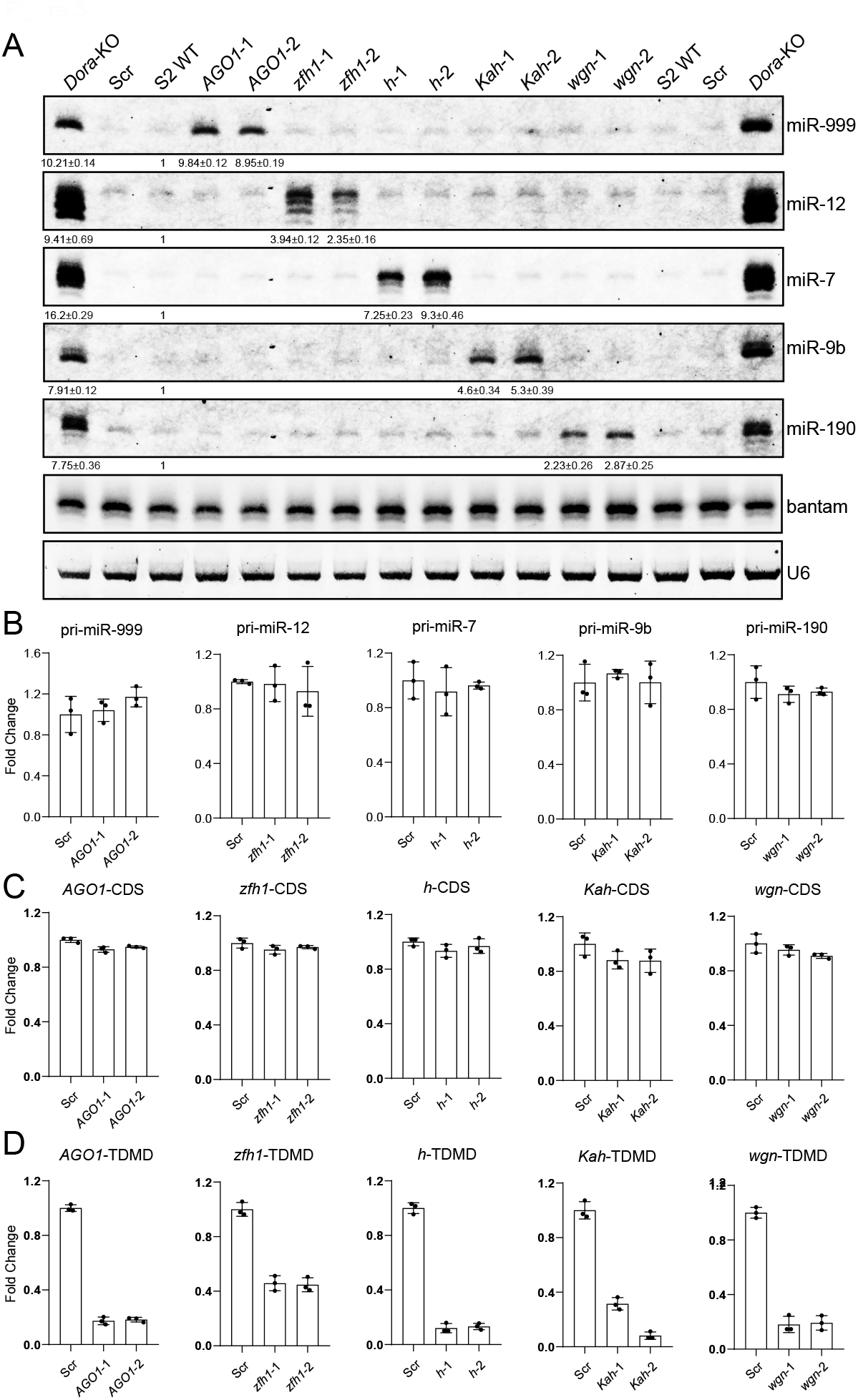
TDMD triggers knockout in S2 cells increase corresponding miRNAs. (**A**) Northern blot analyses of miR-999, miR-12, miR-7, miR-9b, miR-190, bantam and U6 in TDMD trigger knockout of *AGO1, zfh1, h, Kah, wgn* and WT, control-KO (Scramble), *Dora*-KO S2 cells. Total RNAs were extracted from each trigger knockout population cells selected with 5 μg/mL puromycin for 4 weeks. The levels of bantam and U6 serve as loading controls. The miRNA abundance normalized to bantam was shown below miRNA. The miRNA abundance in WT was normalized to 1. n=3 biological replicates. RT-qPCR measurement of the levels of corresponding pri-miRNAs (**B**), CDS region of the trigger transcripts (**C**) and trigger region of the transcripts (**D**) in control and trigger knockout cells.

Increased mature miRNAs by impaired TDMD should not be the result of increased transcription or processing of primary (pri-)miRNAs^14, 15, 16^. To confirm this, we performed RT-qPCR to measure the pri-miRNA levels of the corresponding miRNAs. Our data showed that none of these pri-miRNAs increased in S2 cells after deletion of TDMD triggers, suggesting that the increase in mature miRNAs was not due to increased biogenesis or processing (Figure 3B).

Since tested TDMD triggers are all located in the 3′ UTR of the mRNAs, ranging from 500 to 1300 nt downstream of the stop codon (Figures 2A), we further tested whether knockout of the TDMD triggers affected the expression of the corresponding transcripts. The qPCR results showed that the mRNA levels of the corresponding trigger were not affected after deletion of the TDMD trigger compared to the control (Figure 3C). On the other hand, in each TDMD trigger KO polyclonal cell population, due to the deletion of the TDMD trigger as well as the flanking sequences by CRISPR-Cas9, when we used primers binding near the TDMD trigger for detection, the PCR amplicons were significantly reduced compared with the control (Figure 3D). This further demonstrates that we successfully deleted the TDMD trigger using pairs of sgRNAs targeting each trigger.

We also tested several candidate TDMD triggers which fulfilling two out of the three screening criteria for a high-confidence TDMD trigger, such as *14-3-3epsilon*/miR-277 (missing criteria no. 1, not canonical TDMD pair), *shn*/miR-190 (missing criteria no. 2, hybrid reads <100 RPM) and *CG1673*/miR-277 (missing criteria no. 3, <4 fold hybrid increase in *Dora*-KO) (Figure S3A, Table S1). The results showed that knockout of these TDMD triggers did not affect the corresponding miRNAs (Figure S3B). Taken together, we verified that five endogenous TDMD triggers can induce degradation of the corresponding miRNAs in S2 cells (*AGO1*/miR-999, *zfh1*/miR-12, *h*/miR-7, *Kah*/miR-9b and *wgn*/miR-190).

### TDMD triggers influence miRNA abundance, 3′ end extension and function

To globally verify the loss of TDMD trigger specifically affected the corresponding miRNAs, we performed small RNA sequencing in each trigger KO cells. Consistent with the northern blot results, the abundance of miR-999, miR-12, miR-7 and miR-190 in corresponding trigger KO cells were significantly higher than that in control-KO cells, while other miRNAs had no significant change, including the passenger strands (Figures 4A, 4B, S4A and S4B). In *Kah* trigger KO cells, in addition to the significant increase of miR-9b, miR-996 also increased, suggesting that the *Kah* trigger may potentially regulate other miRNAs (Figure S4C).

**Figure 4.**
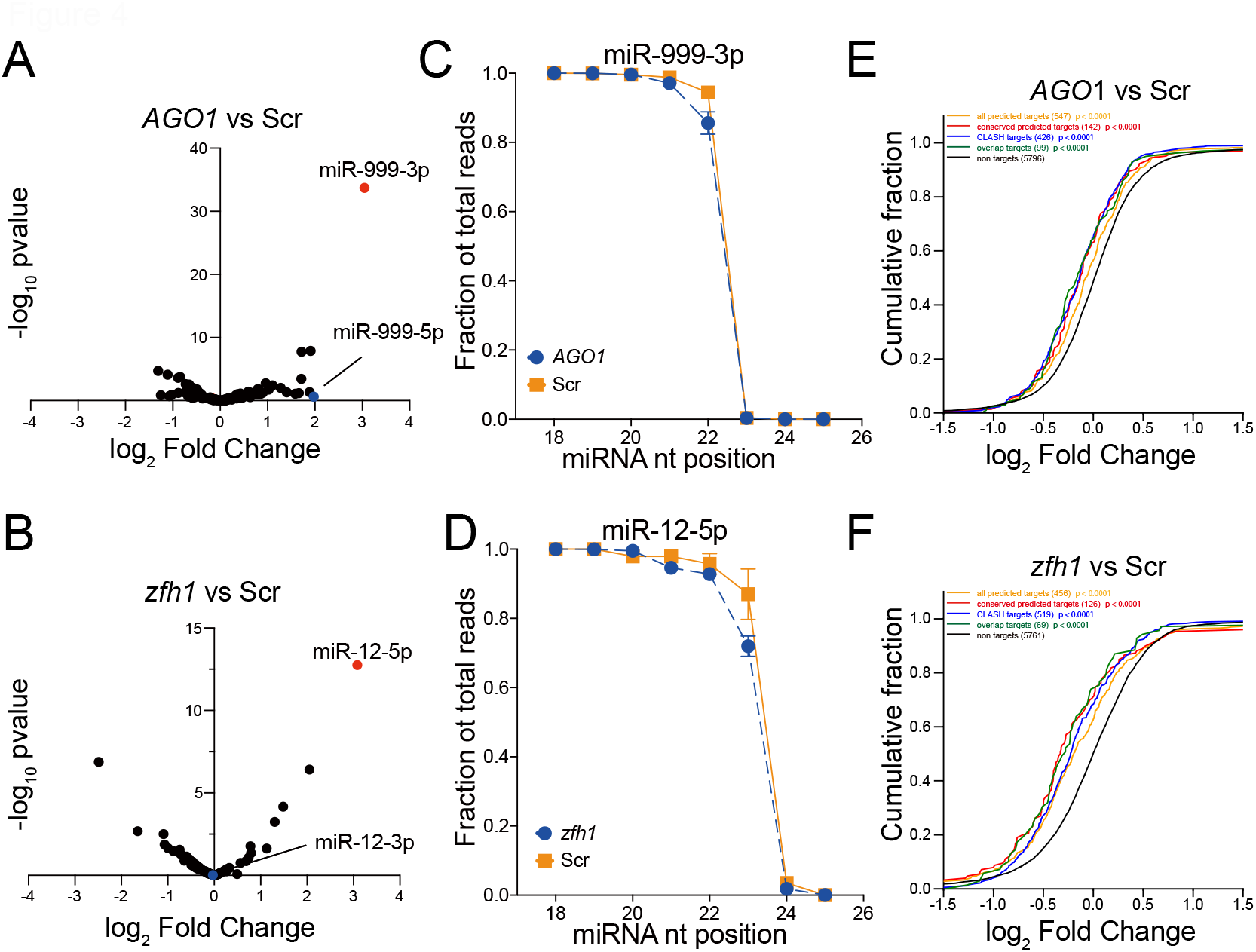
TDMD triggers influence miRNA abundance, 3′ end extension and function. The influence of TDMD trigger on miR-999 (**A**) and miR-12 (**B**) abundance in *AGO1* and *zfh1* trigger-KO cells compared to control-KO cells. miR-999 and miR-12 are indicated by red dots, as the blue dots represent their passenger strands. The fraction of sRNA-seq reads with coverage of 18-26 nucleotides (nt) for miR-999 (**C**) and miR-12 (**D**). For each miRNA, solid lines delineate the control samples, dash lines delineate the TDMD trigger-KO samples. Repression of miR-999 (**E**) or miR-12 (**F**) targets in TDMD trigger-KO cells. Plotted are cumulative distributions of mRNA fold changes observed from TDMD trigger-KO cells, comparing the impact on all predicted miRNA targets (orange line), conserved predicted miRNA targets (red line), CLASH targets (blue line), overlap (CLASH and predicted) targets (green line) to that of non targets (black). Log_2_-fold changes for each set of mRNAs are indicated. Statistical significance of differences between each set of predicted targets and control mRNAs was determined by the Mann–Whitney test.

It is known that the TDMD trigger can also induce miRNA 3′ end tailing and trimming. Therefore, we calculated the numbers of each miRNA from 18 to 26 nt long obtained in the small RNA-seq. As expected, the percentage of miRNAs with 3′ extension was lower when the corresponding TDMD triggers are knocked out (Figures 4C, 4D, and S4D-F).

To examine the functional impact of the TDMD triggers, we performed poly(A) RNA sequencing to detect miR-999 and miR-12 target changes in *AGO1* and *zfh1* trigger-KO cells, respectively. We have five groups of target mRNA including all TargetScan predicted targets, conserved TargetScan predicted targets, CLASH-identified targets, overlap targets (overlap of CLASH and TargetScan predicted targets), and non-targets^28^. Compared with non-targets, the cumulative curves of all four groups of miR-999 and miR-12 targets shift significantly towards the negative direction in *AGO1* and *zfh1* trigger-KO cells (Figures 4E and 4F). Collectively, these data confirmed that deletion of TDMD triggers increases abundance of the corresponding miRNAs in S2 cells, which in turn result in down-regulation of the mRNAs targeted by these miRNAs.

### Morpholinos targeting to TDMD triggers increases miRNA

Out of the five TDMD triggers identified from S2 cells, the *AGO1* trigger is the most conserved (Figure 2A, Table S1), and exhibits the highest efficiency in degrading miR-999 (Figure 3A). Paradoxically, AGO1 protein is essential for global miRNA stability and function, we therefore focus on how the *AGO1* mRNA regulates the abundance of miR-999. To further verify that *AGO1* has a TDMD trigger of miR-999, we designed a 25-base morpholino oligonucleotides (oligos) targeting the TDMD trigger of *AGO1*. Morpholino oligos are synthetic molecules derived from the structure of natural nucleic acids. They bind to complementary sequences of RNA or single-stranded DNA via standard nucleic acid base pairing. Unlike many antisense RNAs (e.g. siRNA), morpholino oligos do not result in the degradation of their target RNA molecules. Instead, morpholino oligos bind to target sequences through “spatial blocking”, and therefore inhibit other molecules from interacting with the target RNA.

After treating S2 cells with 1 µM, 2.5 µM and 5µM *AGO1* or control morpholino oligo for 48 hours, we extracted total RNAs and performed northern blot to detect miR-999 abundance. The abundance of miR-999 gradually increased with increasing concentration of *AGO1* morpholino oligo, which would block the interaction between miR-999 and *AGO1* trigger (Figure 5A). In contrast, there was no change in miR-999 levels when control morpholino oligo was used (Figure 5A). We also performed RT-qPCR to measure the levels of pri-miR-999, *AGO1*-CDS (with primers amplifying a region in the coding sequence) and *AGO1*-TDMD (with primers spanning the TDMD trigger) after morpholino oligo treatment. The results were as expected: since *AGO1* morpholino oligo only blocked miR-999 binding to *AGO1* TDMD trigger, there was no difference in the abundance of pri-miR-999, *AGO1*-CDS and *AGO1*-TDMD between *AGO1* and control morpholino oligo treatments (Figure 5B). These experiments further confirmed the TDMD interaction between *AGO1* and miR-999.

**Figure 5.**
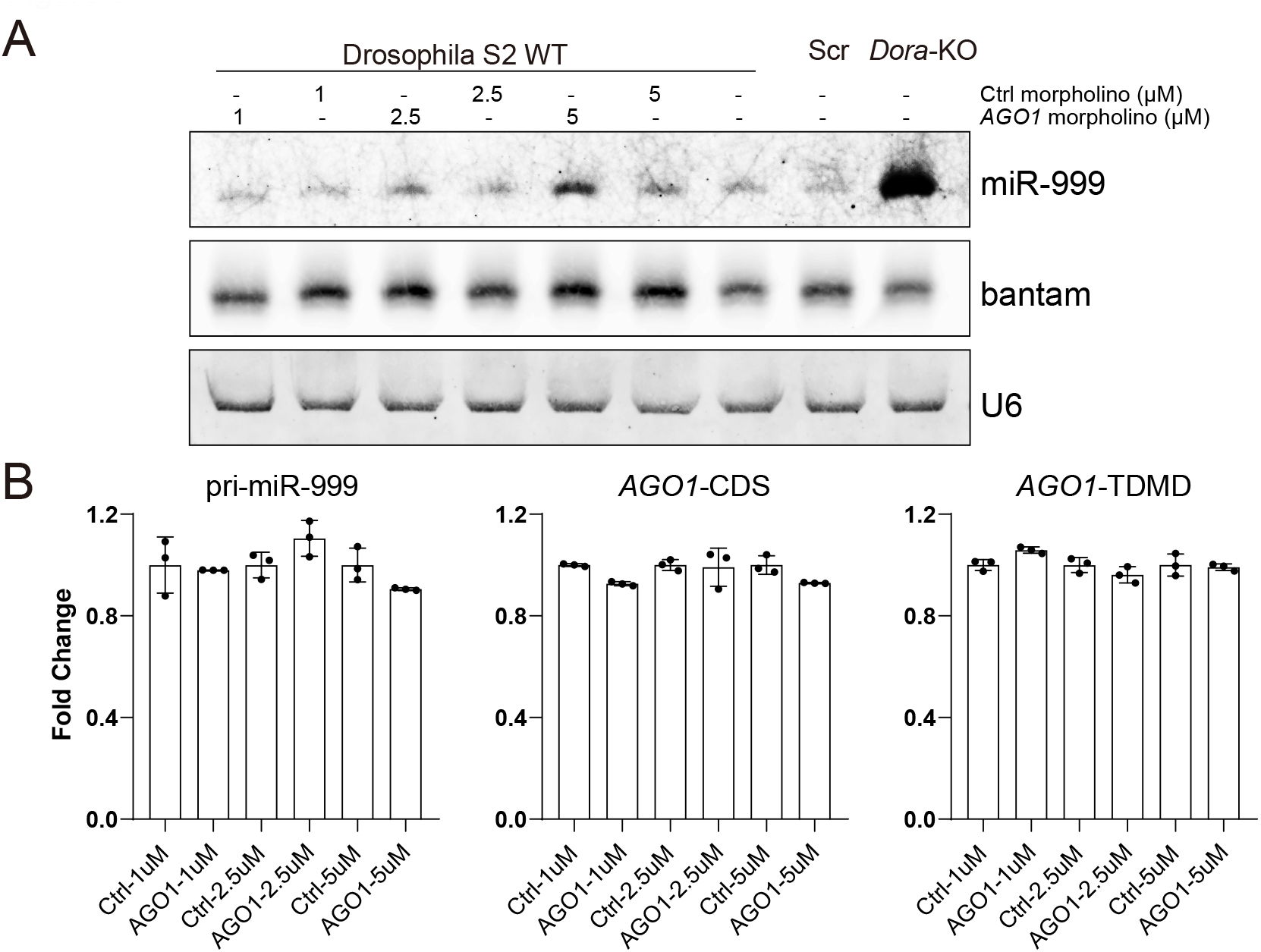
Morpholino oligos targeting *AGO1* trigger increases miR-999. (**A**) Northern blot analyses of miR-999, bantam and U6 in S2 cells following 48 h of 1µM, 2.5µM or 5µM morpholino oligos treatment. Bantam and U6 served as loading controls. (**B**) RT-qPCR analyses of pri-miR-999, *AGO1*-CDS and *AGO1*-trigger levels in control and *AGO1* morpholino oligos treated cells as in A.

### Deletion of *AGO1* trigger increases miR-999 in *Drosophila*

To further investigate the biological function of *AGO1* trigger in *Drosophila*, we constructed mutant strains with the *AGO1* trigger deleted by CRISPR-Cas9 (Figure 6A). Similar to the CRISPR KO performed in S2 cells, we used two sgRNAs targeting both sides of the *AGO1* trigger. Two sgRNAs used for *AGO1* trigger KO in S2 cells were cloned into the pCFD6 which can liberate multiple functional sgRNAs from a single precursor transcript by a tRNA–sgRNA expression system^29^. After injecting the plasmid to embryo, transgenic flies with a pCFD6 construct containing two sgRNAs targeting the *AGO1* trigger (pUAS-tRNA-sgRNA1-tRNA-sgRNA2-tRNA) were crossed to flies expressing Cas9 (pUAS-Cas9, nos-GAL4::VP16) (Figure 6A first cross). Subsequently, we obtained virgin female flies to cross with *Cyo* balancer flies twice to obtain the potential lines containing *AGO1* trigger deletions (Figure 6A second and third crosses). This approach generated 11 lines containing different deletions by genotypic identification (Figure 6B).

**Figure 6.**
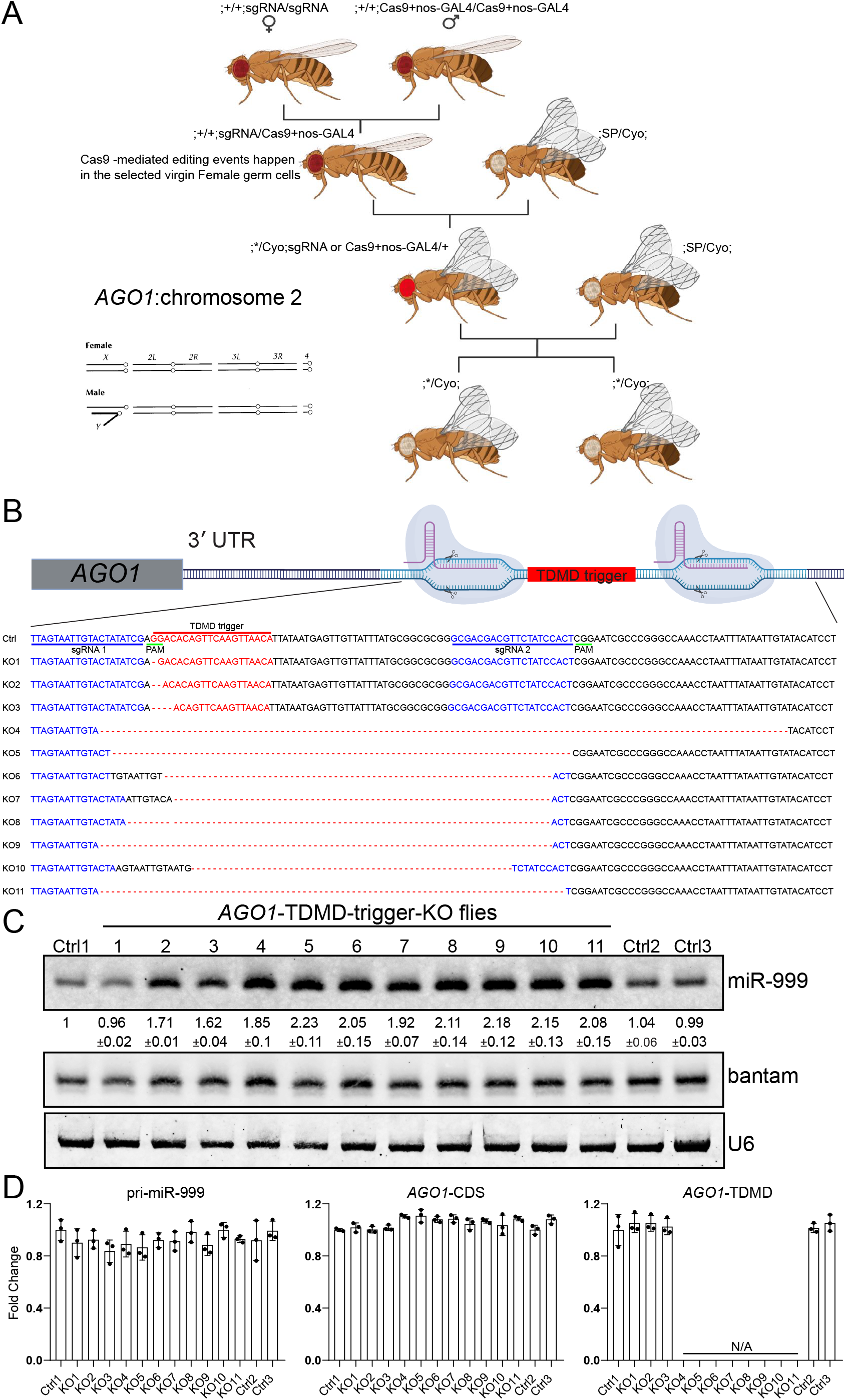
Deletion of *AGO1* trigger increases miR-999 in *Drosophila*. (**A**) Schematic showing the generation of *AGO1* trigger KO flies. Virgin female flies containing *AGO1* trigger-targeting CRISPR-Cas9 in the germ cells were crossed twice with Cyo balancer flies to obtain potential lines containing the *AGO1* trigger deletion.*represents mutation of the *AGO1* trigger. (**B**) Schematic of the CRISPR-Cas9-mediated knockout of miR-999 TDMD trigger from *AGO1* 3′ UTR. The trigger and sgRNAs are highlighted in red and blue, respectively. PAM sequence is underlined (green). The genotype of each mutant line is shown below. (**C**) Northern blot analyses of miR-999, bantam and U6 in controls and *AGO1* trigger mutant lines. Bantam and U6 served as loading controls. The normalized miR-999 abundance (compared to bantam) are shown below each miRNA band. The miRNA abundance in control line no.1 was normalized as 1. n=3 biological replicates. Control lines without mutation were obtained from same crossing procedure as the mutant lines. (**D**) RT-qPCR analyses of pri-miR-999, *AGO1*-CDS and *AGO1* trigger levels in controls and *AGO1* trigger mutant lines as in C.

We next tested miR-999 abundance in various deletion mutant *Drosophila* lines by norther blot. We found that the abundance of miR-999 doubled in KO lines with complete deletion of the *AGO1* trigger (KO4-KO11) compared to control flies (Figure 6C), which were obtained through the same injection and crossing procedure as the KO lines but has been sequence-verified as having no mutation at or near the *AGO1* trigger. In the partial deletion mutant lines, we found that when only the last nucleotide (G) base-paired with the 3′ end of miR-999 (U) was missing (KO1), the mutant did not show increased abundance of miR-999. This is probably because the A after the deleted G can still base-pair with the last nucleotide of miR-999. However, when the deletion increased to two or four bases (KO2 and KO3), the abundance of miR-999 increased to comparable levels as the complete trigger KO lines (Figure 6C).

Since miR-999 is known to be highly expressed in the head, particularly in the brain according to the FlyAtlas2 database, we also examined the changes of miR-999 in the head and body of the flies separately. Indeed, miR-999 is highly expressed in the head and are barely detectable in the body (Figure S5). However, consistent with the result obtained from the whole fly, the abundance of miR-999 in both body parts from all *AGO1* trigger-KO lines significantly increased compared with that in control flies, except for KO1 (Figure. S5). These results further suggest that the pairing of trigger with miRNA 3′ end is crucial for TDMD.

In addition, the expression levels of pri-miR-999, *AGO1* coding region and TDMD trigger region were also detected by RT-qPCR. The results showed that the levels of pri-miR-999 and *AGO1* coding region were not changed by knockout (Figure. 6D). The unchanged *AGO1* mRNA levels suggest that trigger transcripts are not under miRNA-mediated repression, consistent with the recent findings^26^. In contrast, because KO4 to KO11 do not contain the TDMD trigger and flanking sequences, no signal can be detected by the primer spanning the TDMD trigger. Because knockout lines KO1 to KO3 contain only up to 4 nt deletions, PCR amplicon of their TDMD region remain unchanged (Figure 6D). Therefore, *AGO1* trigger modulates the levels of miR-999 not only in S2 cells but also in adult flies.

### *AGO1* trigger-KO flies are more vulnerable to oxidative stress

Information on the FlyBase indicates that overexpression of miR-999 in fly resulted in lethality^30^, implying that miR-999 plays an important biological function in *Drosophila* development. However, homozygous *AGO1* trigger-KO fly was not lethal, probably because the abundance of miR-999 increased only about 2-fold (Figure 6C). To further investigate the biological effect of *AGO1* TDMD trigger on flies, we performed small RNA and poly(A) RNA sequencing in trigger deletion *Drosophila* lines. Consistent with the results of the northern blots, the abundance of miR-999 in selected *AGO1* trigger KO lines (KO9-KO11) is 2-fold higher than control flies, while other miRNAs had no significant change, including the passenger strand of miR-999 (Figure 7A). Meanwhile, miR-999 target RNAs were significantly downregulated in mutant lines in poly(A) RNA sequencing (Figure 7B).

**Figure 7.**
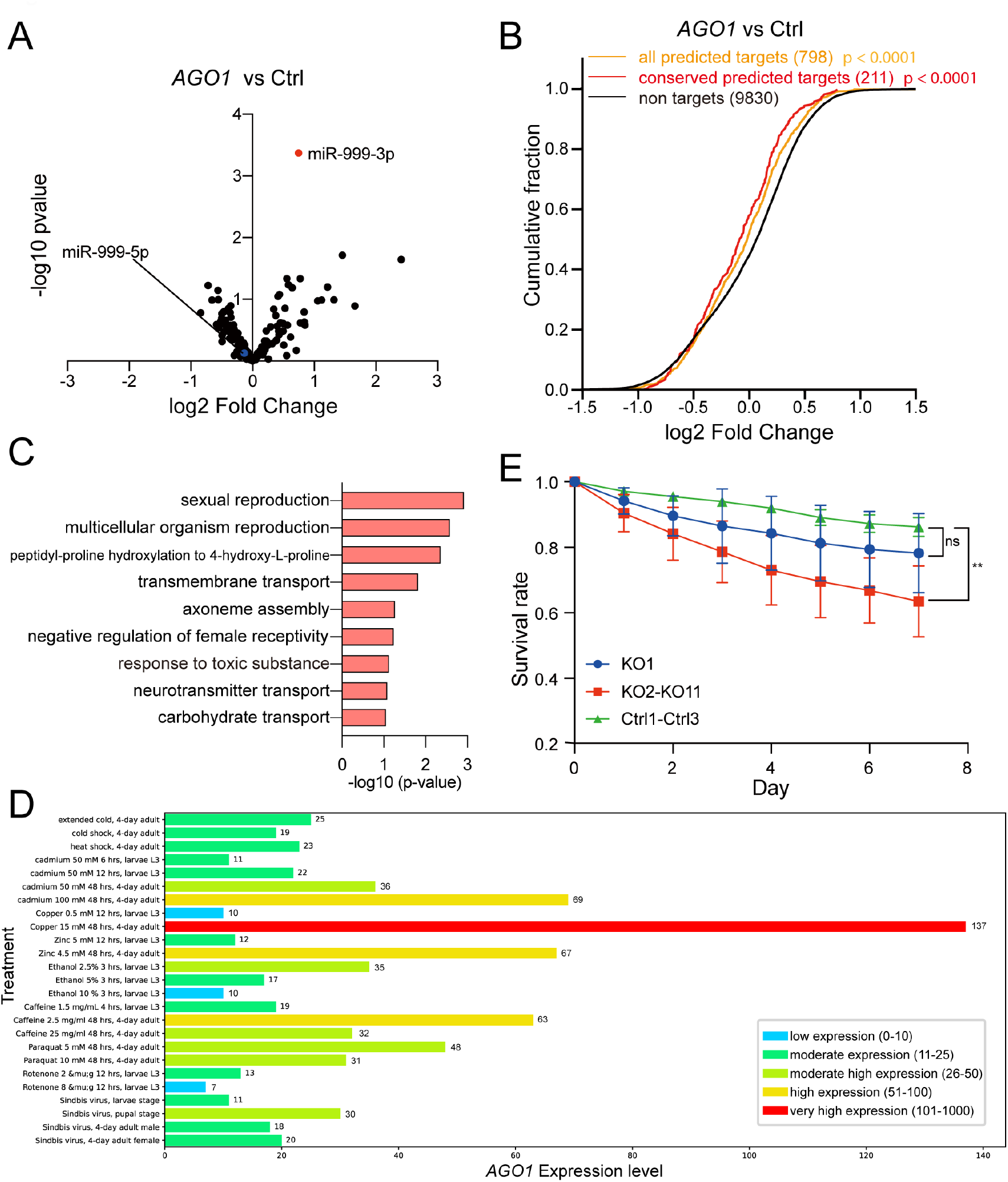
AGO1 trigger-KO flies are more vulnerable to stress. The influence of TDMD trigger on miR-999 (**A**) and repression of miR-999 targets (**B**) in *AGO1* trigger KO flies. (**C**) DAVID identified GO term biological pathways enriched in down-regulated genes in *AGO1* trigger KO flies compared with control-KO flies. (**D**) The expressions of *AGO1* after different treatments were generated by modENCODE of FlyBase. (**E**) Survival rate for *AGO1* trigger KO flies after hydrogen peroxide exposure compared with control-KO flies. (∗*) P< 0.01, paired t test. Four independent oxidative stress experiments were conducted. In each experiment, each fly line has three vials starting with 20 male flies.

Since the *AGO1* trigger-KO flies do not exhibit apparent lethality phenotype, we tested partial lethality as well as lifespan but there were no significant differences in mutant lines compared with control flies (data not shown). Based on gene ontology (GO) term enrichment analyses of the significantly down-regulated genes in the *AGO1* trigger-KO flies and S2 cells, miR-999 may be involved in several stress response pathways including response to toxic substance, response to wounding and defense response to bacteria/fugus (Figures 7C and S6). Accordingly, *AGO1* expression is strongly induced in response to stresses such as high concentration of copper, which is known to induce oxidative stress (Figure 7D)^31, 32^. Therefore, we investigated the effects of *AGO1* TDMD trigger KO on the lifespan of *Drosophila* when exposed to hydrogen peroxide, a reactive oxygen species that are routinely used to induce oxidative stress in flies^33, 34^. We fed 9-12 day old flies with 1% sucrose solution containing 4M hydrogen peroxide for 24 hours and returned them to normal food. The flies were then counted daily to monitor survival. After one week, the survival rate of mutant lines with effective *AGO1* trigger KO, but not KO1, was significantly lower than that of the control flies (Figure 7E). These data suggest that the *AGO1* TDMD trigger is required for optimal stress response via inducing miR-999 degradation.

## Discussion

As post-transcriptional regulators, the regulation of miRNA biogenesis has been extensively studied, while the mechanism of its degradation is less well understood. TDMD initiates miRNA degradation through extended pairing between miRNA and target RNA, which appears to be the main mechanism regulating miRNA turnover^4, 5^. Previous studies have identified several representative TDMD triggers in mammalian cells, as well as the most critical enzyme in the TDMD pathway, ZSWIM8^6, 7, 13, 14, 15, 16^. Many *Dora*-sensitive miRNAs have been found in *Drosophila*, which potentially subject to TDMD degradation mechanisms^7, 12^. A recent report described the identification of six TDMD triggers from *Drosophila* S2 cells and embryos via bioinformatic prediction, which predicted about a dozen triggers for experimental validation^26^. In this study, we performed a large-scale screening of TDMD triggers for sensitive miRNAs in *Dora*-KO S2 cells by AGO1-CLASH. By setting strict screening criteria of TDMD base-pairing pattern, high RNA/miRNA hybrid abundance (>100 RPM) and enrichment of the hybrid in *Dora*-KO, we efficiently identified five endogenous triggers that can degrade the corresponding base-paired miRNAs, including *AGO1*/miR-999, *zfh1*/miR-12, *h*/miR-7, *Kah*/miR-9b and *wgn*/miR-190.

Compared to our previous TDMD trigger identification using AGO-CLASH in mammalian cells without perturbation of the TDMD pathway^13^, here we performed *AGO1*-CLASH in *Dora*-KO cells. Since Dora is a key enzyme in the TDMD pathway, deletion of *Dora* greatly increases the abundance of miRNAs regulated by TDMD and presumably stabilizes the interactions of TDMD miRNA/target RNA hybrids, which facilitates the identification of TDMD triggers with high confidence. Therefore, performing AGO-CLASH in the *ZSWIM8* null background can identify genuine TDMD triggers more confidently.

Interestingly, based on our CLASH screen, *Kah* may act as a trigger for both miR-9b and miR-9c. Subsequent *Kah* trigger-KO experiments revealed that this trigger only affects the abundance of miR-9b, while miR-9c remained unchanged (Figures 3A and S2). Further analysis showed that the base-pairing pattern between miR-9c and *Kah* is more extensive at the miRNA 3′ end, while miR-9b and *Kah* have two mismatches at the 3′ end (Figure 1D). Moreover, hybrid reads of *Kah*/miR-9c are about twice as many as *Kah*/miR-9b in *Dora*-KO *AGO1*-CLASH (Table S1). These analyses would suggest that *Kah* is more likely to be a trigger of miR-9c than miR-9b. However, the *Kah*/miR-9b does have one more base-pairing at nt 9 of the miRNA compared with *Kah*/miR-9c, suggesting that this base pair is critical in contributing to the higher TDMD efficiency for *Kah* trigger towards miR-9b than miR-9c.

The five TDMD triggers we found are all in the 3′ UTR, and we also tested triggers located in the CDS region such as *shn* trigger. The base-pairing pattern for *shn*/miR-190 is highly extensive, with 10 base pairs extended to the very 3′ end of miRNA (Fig. S3A). However, the abundance of miR-190 was not affected when *shn* trigger was knocked out (Figure S3B). Meanwhile, all known mammalian triggers are in the 3′ UTR of mRNA or noncoding RNAs, probably because the interactions of miRNA and CDS region triggers are crowded out by the translating ribosomes.

*Dora*-sensitive miRNAs showed completely different compositions in S2 cells and embryos. miR-7, miR-9b, miR-9c, miR-12, miR-190, and miR-999 were significantly elevated in *Dora*-KO S2 cells (Figure 1). On the other hand, miR-3 and miR-310 family members were significantly increased in *Dora*-KO embryos (Figure S1). Similarly, when the *AGO1* trigger was knocked out, the magnitude of the miR-999 increase was much weaker in flies than in S2 cells (Figures 3 and 6). Also, *h* and *Kah* are effective triggers for miR-7 and miR-9b in S2 cells, respectively, but when we knocked out these two triggers in *Drosophila*, we were surprised to find that they did not affect the corresponding miRNAs in the total RNA extracted from adult flies (Figure S7). These results indicate that TDMD are specific in different types of cells and tissues, and different stages of development.

In this study, we found that *AGO1* trigger can induce miR-999 degradation in both S2 cells and *Drosophila*. Interestingly, in *Drosophila*, miRNAs are normally loaded onto the AGO1 protein to form a complex, thus the miRNAs are protected from degradation. However, the TDMD trigger located in the 3′ UTR of *AGO1* can specifically degrade miR-999. Therefore, the *AGO1* coding region and the TDMD trigger formed a bidirectional regulation of the abundance of miR-999 at the protein and RNA levels, respectively. Interestingly, there are four alternative forms of *AGO1* 3′ UTR due to alternative polyadenylation, with the shortest 3′ UTR lacking the TDMD trigger (Figure 2A). Future studies are required to determine whether there exists a regulation mechanism by alternative polyadenylation to express *AGO1* mRNAs with different forms of 3′ UTR, which may specifically control miR-999 levels in different cells.

## Materials and Methods

### Cell culture

*Drosophila* S2 cells were cultured at 28°C in Schneider’s Insect Medium (Sigma) supplemented with 10% heat-inactivated bovine growth serum (HyClone). When confluent, the S2 cells were passaged with 1:5 dilution every 5 days. For transfections, 2.5×10^6^ S2 cells/well were seeded in 6-well plates for 24 hours, and then transfected using lipofectamine 3000 (Invitrogen) according to the manufacturer’s protocols. For morpholino oligo treatment, 6×10^6^ S2 cells/well were seeded in 6-well plates and incubated with 1µM, 2.5µM or 5µM vivo-morpholino oligos against *AGO1* TDMD trigger in the medium for 48 hours, then the total RNA was extracted for northern blot and RT-qPCR. Morpholino oligo sequences and RT-qPCR primers are listed in Table S2.

### AGO1-CLASH

AGO1-CLASH was performed in *Drosophila* S2 cells, with *Dora*-KO or control-KO as previously described with minor modifications^35^. Cells were washed in ice-cold PBS, and irradiated with 254 nm UV at 400 mJ/cm^2^. Cell pellets were snap frozen in liquid nitrogen and stored at -80°C until lysis. Pellets were thawed on ice and lysed in lysis buffer (50 mM Tris– HCl pH 7.4, 100 mM NaCl, 1% NP40, 0.1% SDS, 0.5% sodium deoxycholate, Complete Protease Inhibitor to a final concentration of 2 ×, Murine Inhibitor (NEB) 40 U/ ml lysis buffer; 1 ml lysis buffer per approximately 0.3 × 10^9^ cells, 1.2 × 10^9^ cells per sample) for 15 minutes, then treated with 20U/mL RQ1 DNase (Promega) for 5 minutes at 37°C with shaking, and centrifuged at 21,000xg for 15 minutes at 4°C. AGO1-IP was carried out with magnetic protein A dynabeads (Life Technologies) (100 µl) conjugated with polyclonal AGO1-antibody (Abcam #ab5070, 20 µg per sample) and incubated with cell lysate overnight at 4°C as previously described^36^. After IP, samples were washed three times with lysis buffer and treated with 15 ng/μL RNaseA for 12 minutes at 22°C. After that, intermolecular ligation and libraries were generated as previously described^35^. Libraries were separated and size selected between 147 and 527 bp as previously described^37^. Three AGO1-CLASH libraries were generated from control-KO and *Dora*-KO S2 cells respectively. Libraries were sequenced on the Illumina NovaSeq 6000 by the University of Florida Interdisciplinary Center for Biotechnology Research (ICBR) NextGen DNA Sequencing Core.

### Identification of TDMD triggers in S2 cells CLASH data

The adapter sequences of raw reads were trimmed with Cutadapt software^38^, and reads < 18 nt were removed. Paired FastQ files were assembled by Pear software^39^ and collapsed by fastx_collapser (http://hannonlab.cshl.edu/fastx_toolkit) based on the 4 random nucleotides at the 5′ end and 3′ end of the reads to remove PCR duplications. Then random nucleotides were removed by Cutadapt software. Adapter sequences are listed in Supplemental Table S2. Processed reads then underwent mapping, hybrid calling, base-pairing prediction, and annotation by the Hyb software with default settings^40^. Note that in the default settings, the target RNA reads were extended 25 nt on the 3′ end to compensate for possible trimming of the sequences that can base-pair with the miRNA.

Custom Python scripts (“python3 CLASH.py TDMD_analyzer -i [--input]” in https://github.com/UF-Xie-Lab/TDMD) were written to screen candidate TDMD hybrids that satisfied four criteria based on reported TDMD pairs: First, the seed region (nucleotides 2–8) of the miRNA can base-pair with the target RNA, allowing G-U wobble pairs. Second, the 3′ end of the miRNA must contain more than 7 consecutive base pairs with the target RNA within the last 8 nucleotides, or contain 9 consecutive base pairs. Third, the central bulge of the target RNA/miRNA hybrids should be <7 nt, but >0 nt. Four, the binding energy between miRNA and the target should be lower than 16 kcal/mol. TDMD hybrids that met these criteria are summarized in Supplemental Table S1.

### RNA-seq analysis

*AGO1* trigger KO Small RNAs (18-40 nts) and poly-A selected RNA from S2 cells were sequenced in Novogene, other small RNAs were sequenced by BGI. All sequencing experiments were performed in duplicates. For small RNA-seq reads from Novogene, the adapters were removed by Cutadapt software (version 3.4); for clean-miRNA-seq reads from BGI, the reads were collapsed based on the unique molecular identifiers (UMIs) using custom python scripts (“python3 CLASH.py deduplicate_BGI -i [--input] <fastq>“). For clean miRNA abundance and length distribution analysis, we calculated clean reads that can match the 18 nts of the annotated miRNAs using custom python scripts (“python3 CLASH.py miRNA_abundance -i [--input] <fasta/fastq> -d [--miRNA_database]” (for abundance analysis) or “python3 CLASH.py miRNA_length_distribution -i [--input] <fasta/fastq> -d [--miRNA_database]]” (for length distribution analysis) on https://github.com/UF-Xie-Lab/TDMD).

For poly-A RNA-seq reads, the adapters were removed by Cutadapt software (version 3.4), and then the clean FASTQ file was mapped to the *Drosophila melanogaster* genome (FlyBase Release 6.32) by Hisat2 (version 2.2.1) software. Subsequently, we used HTSeq-count software (version 0.11.1) to count each gene’s abundance. The differential expression levels of each miRNA or poly-A RNA were calculated using the Deseq2 package (version 1.20.0) of R, the miRNA normalization was performed by Deseq() function. miRNAs with baseMean (the average of normalized count values) below 200 and mRNAs with baseMean below 100 were filtered out. The cumulative fraction curves (CFC) were drawn by matplotlib (version 3.4.1) using a custom python script (“python3 CLASH.py Cumulative_fraction_curve_targetScan_CLASH -i [--DEseq_file] -a [--all_targets] -c [--conserved_targets] -l [--clash_targets] -t [--clash_targetScan_interacted] -b [--baseMean]” in https://github.com/UF-Xie-Lab/TDMD).

### Gene Expression Profile Analyses

For tissue-specific expression of miRNA and genes, genome-wide tissue-specific gene expression profiles from the FlyAtlas 2 database were used. FlyAtlas 2 is a repository of tissue-specific gene expression based on genome-wide RNA-seq analyses of *D. melanogaster* genes in adult males, females, and larvae. Pearson correlation analysis was calculated by Prism 8.

### GO Term enrichment analysis

From the poly-A RNA-seq data, 167 down-regulated genes in *AGO1* trigger KO S2 cells and 93 down-regulated genes in *AGO1* trigger KO flies (baseMean>100; log FoldChange < -1, p value<0.05) were analyzed by DAVID Bioinformatics Resource v2022q3 (https://david.ncifcrf.gov)^41, 42^. The top 10 biological processes from respective data sets are graphed according to the -log10(p-value) in Prism 8.

### Plasmid construction

For knockout of the candidate TDMD triggers, three or four different sgRNAs for each trigger were designed and cloned into the pAc-sgRNA-Cas9 (Addgene #49330) vector^27^. For knockout of TDMD trigger in flies, two sgRNAs were cloned into the pCFD6 (Addgene #73915) vector which is a tRNA–sgRNA expression system using the endogenous tRNA processing machinery liberating multiple functional sgRNAs from a single precursor transcript in the nucleus^29^, sgRNA (guide sequences) are listed in Table S2.

### Knockout cell lines

To generate TDMD trigger knockout cell lines, two different sgRNA plasmids targeting both sides of the TDMD trigger were co-transfected into S2 cell by lipofectamine 3000 (Invitrogen) according to the manufacturer’s protocols, sgRNA (guide sequences) are listed in Table S2. 72 hours after transfection, cells were selected with 5 μg/mL puromycin (Life Technologies) for 4 weeks.

### Fly transgenesis and culture

pCFD6 construct containing two sgRNAs targeting the TDMD trigger (pUAS-tRNA-sgRNA1-tRNA-sgRNA2-tRNA) were injected into embryo to generate transgenic lines using standard PhiC31-integrase–mediated transformation to the second (*h* and *Kah* knockout) or third (*AGO1* knockout) chromosome by Rainbow Transgenic Flies. sgRNA guide sequences are listed in Table S2. Flies expressing Cas9 (pUAS-Cas9, nos-GAL4::VP16) are available from the Bloomington *Drosophila* Stock Center (stock number 54593). To generate TDMD trigger knockout and control lines, Virgin females expressing transgenic sgRNAs were crossed to males transgenic for Cas9. Subsequently, the obtained virgin female flies were crossed with CyO or TM3 balancer flies twice to obtain the potential lines containing TDMD trigger deletions. All crosses were performed at 25 °C with 50 ± 5% relative humidity and a 12-h-light– 12-h-dark cycle.

### Northern blot

Northern blots were performed as previously described with infrared probe and EDC (N-(3-dimethylaminopropyl)-N′-ethylcarbodiimide) (Thermo Scientific) crosslinking^43, 44^. Probes are listed in Table S2. Total RNA was extracted with Trizol Reagent (Life Technologies) according to the manufacturer’s protocol and separated on 15% Urea PAGE, then transferred to Hybond-NX membrane (GE Healthcare). The results were analyzed with ImageQuant TL (v7.0).

### Oxidative Stress Assays

Within the 72 hours after initial eclosion, males were separated from the females and held in normal food vials for 8 days, making them 9-12 days old. At this age, the flies were then exposed to 4M hydrogen peroxide 1% sucrose solutions for 24 hours in empty vials. These vials were prepared by adding 2mL of the 4M hydrogen peroxide 1% sucrose solutions to folded KimWipes filter paper. After exposure for 24 hours, the flies were anesthetized by CO_2_ and moved to vials containing normal fly food at about 20 male flies per vial. Finally, their lifespans were assayed daily.

## Data availability

All custom scripts have been made available at https://github.com/UF-Xie-Lab/TDMD. Additional modified scripts can be accessed upon request. All sequencing data that support the findings of this study have been deposited in the National Center for Biotechnology Information Sequence Read Archive (SRA) and are accessible through the BioProject PRJNA896239. All other relevant data are available from the corresponding author on request.

## Acknowledgements

We thank Dr. Elena Kingston and Dr. David P. Bartel for kindly providing *Dora*-KO S2 cell; Dianshu Zhao and Dr. Adam Wong for assisting with *Drosophila* tests. This work was supported by grants from National Institutes of Health (R35GM128753 to M.X.), American Cancer Society (Research Scholar Award to M.X.) and Florida Department of Health (Live Like Bella Pediatric Cancer initiative 21L03 to M.X.).

## Authors Contributions

P.S., L.L. and M.X. conceived the project and were involved in all experiments. T.L., Y.W. and N.M.H. measured RNA levels by RT-qPCR, extracted genomic DNA from flies for genotyping. J.S.M. and J.S.S. collected *Dora* mutant embryo, L.Z. supported fly culture and generation of mutants. P.S., L.L. and M.X. wrote the paper, with comments from L.Z..

---

**Table S1. List of candidates TDMD hybrids in *Drosophila* S2 cells**.

**Table S2. List of adapter for sequencing library processing and oligonucleotides for plasmid construction, PCR, northern blot and morpholino**.

**Figure S1.**
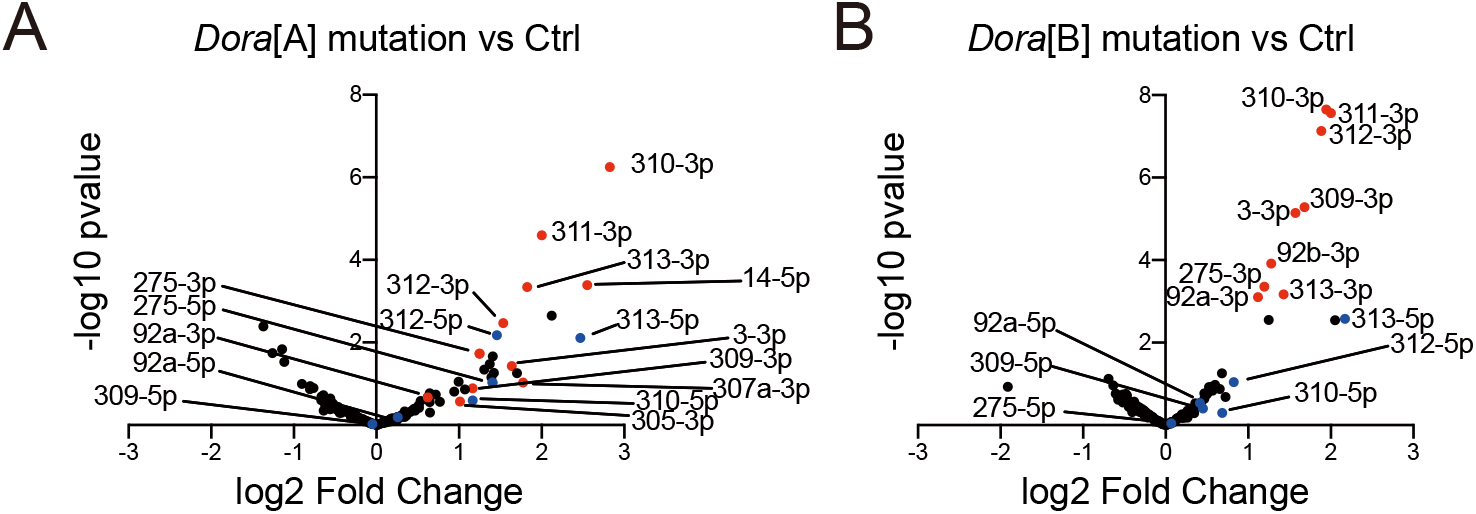
miRNA abundance in embryos with *Dora* mutations. Changes in miRNA abundance observed from *Dora*[A] (**A**) or *Dora*[B] (**B**) embryos compared with control embryos by small RNA-seq. Guide strands of the Dora-sensitive miRNAs are indicated by red dots, and the blue dots represent their passenger strands.

**Figure S2.**
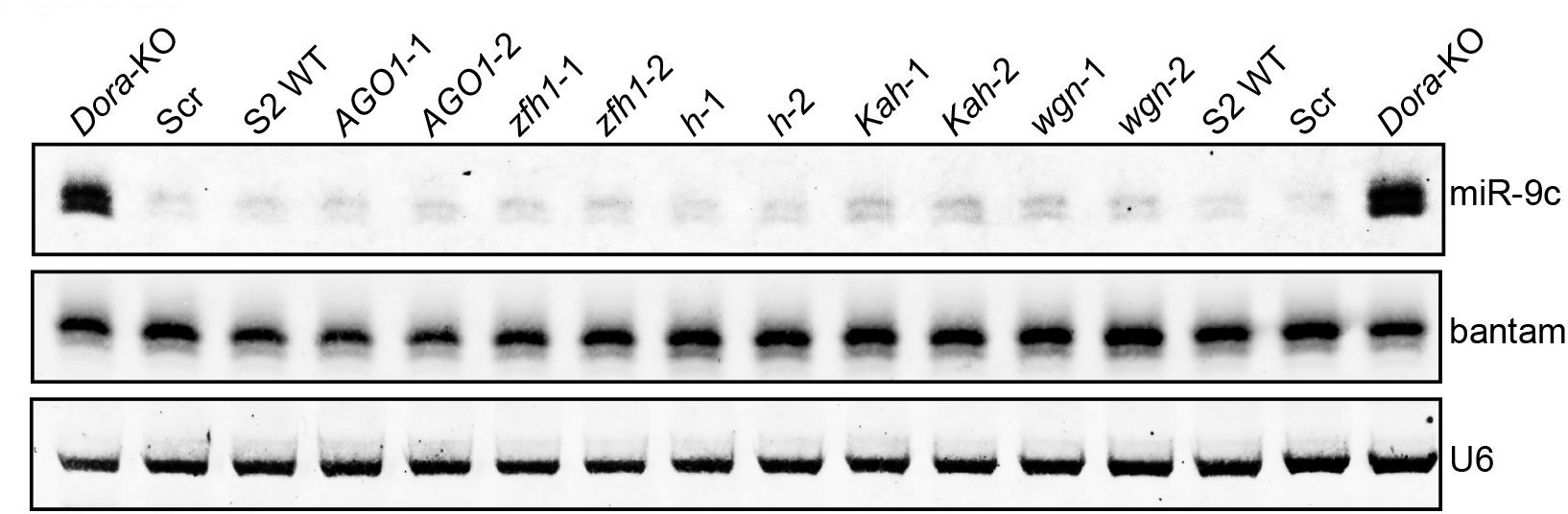
Northern blot analyses of miR-9c in TDMD trigger knockout cells. Northern blots detecting miR-9c, bantam and U6 in TDMD trigger knockout of *AGO1, zfh1, h, Kah, wgn* and WT, control-KO (Scramble), *Dora* KO S2 cells. Total RNAs were prepared as described in Figure. 3A.

**Figure S3.**
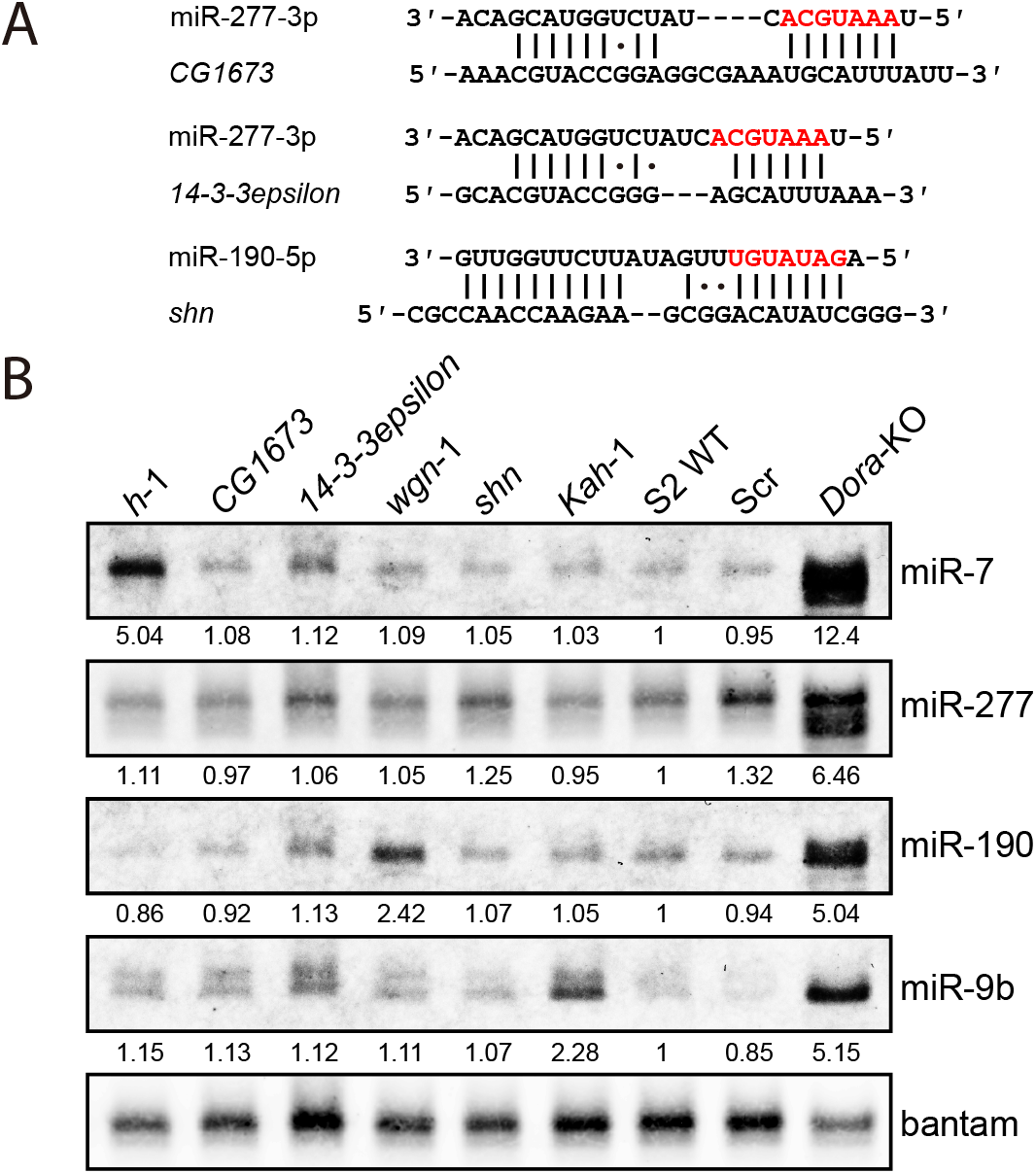
Knockout low-confidence TDMD trigger in S2 cells. (**A**) Base-pairing pattern of miRNAs and potential TDMD triggers. Red letters represent miRNA seed region. (**B**) Northern blot analyses of miR-7, miR-277, miR-190, miR-9b, bantam and U6 in TDMD trigger knockout of *h, CG1673, 14-3-3epsilon, wgn, shn, Kah* and WT, control-KO (Scramble), *Dora*-KO S2 cells. Total RNAs were extracted from each TDMD trigger knockout population cells selected with 5 μg/mL puromycin for 4 weeks. The levels of bantam serve as a loading control. *h*/miR-7, *Kah*/miR-9b and *wgn*/miR-190 serve as positive TDMD pairs. The miRNA abundance normalized to bantam was shown below each miRNA. The miRNA abundance in WT was normalized to 1.

**Figure S4.**
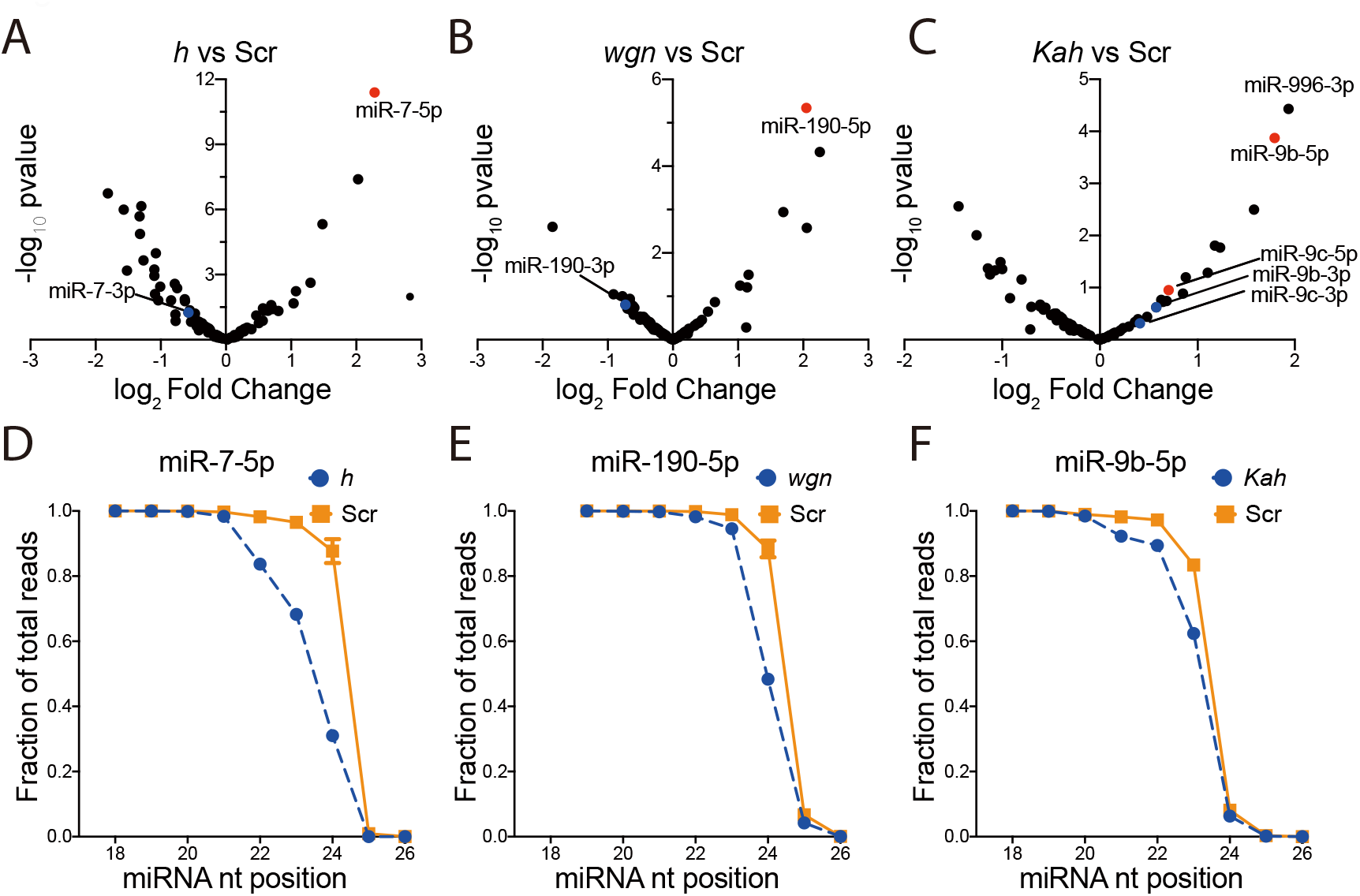
TDMD triggers influence miRNA abundance, 3′ end extension. The miRNA abundance change detected by small RNA-seq in *h* (**A**), *wgn* (**B**) and *Kah* (**C**) TDMD trigger KO cells compared with control-KO (Scramble) cells. miR-7, miR-190 and miR-9b are indicated by red dots, and the blue dots represent their passenger strands. The fraction of small RNA-seq reads with coverage of 18-26 nucleotides (nt) for miR-7 (**D**), miR-190 (**E**) and miR-9b (**F**). For each miRNA, solid lines delineate the control KO samples, dash lines delineate the TDMD trigger KO samples.

**Figure S5.**
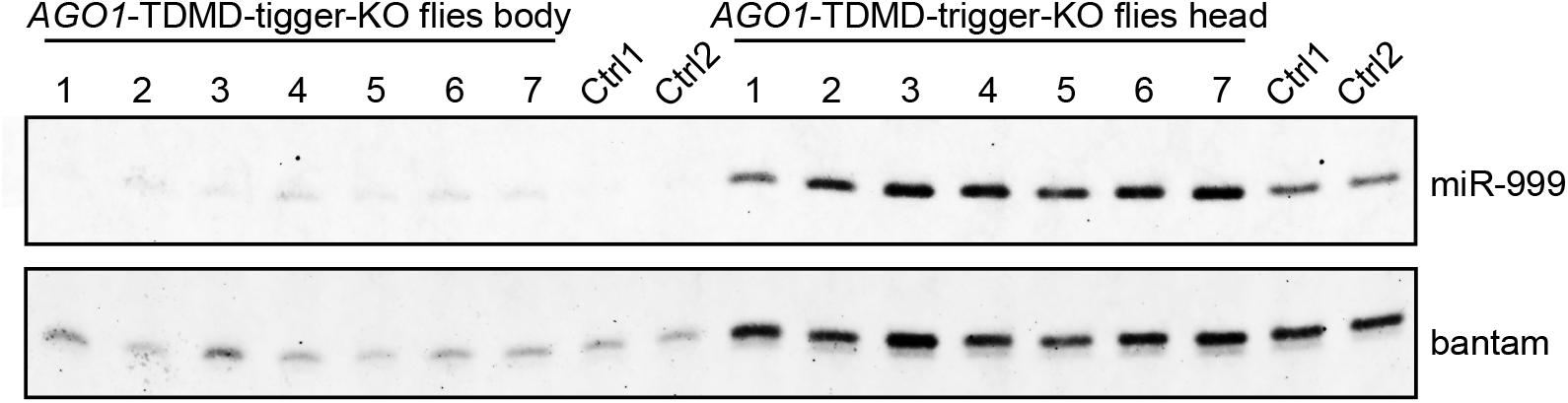
Northern blot analyses of miR-999 in *AGO1* trigger deletion flies. Total RNAs were extracted from the head and body of 30 flies. Samples of head and body were loaded at 20 and 30 ug total RNA, respectively. Bantam serves as a loading control.

**Figure S6.**
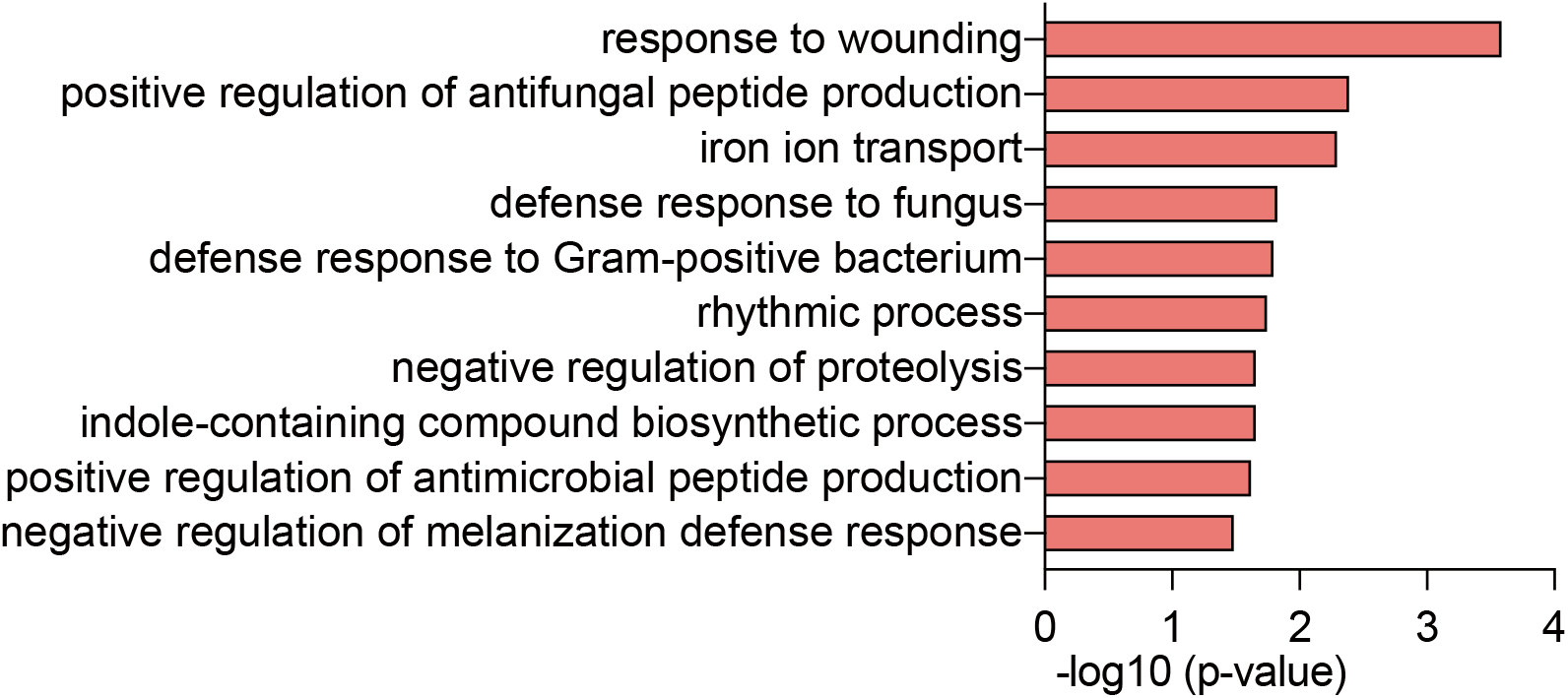
Biological functions of down-regulated genes in *AGO1* trigger KO S2 cells. DAVID identified GO term biological pathways enriched in down-regulated genes in *AGO1* trigger-KO compared with control-KO S2 cells.

**Figure S7.**
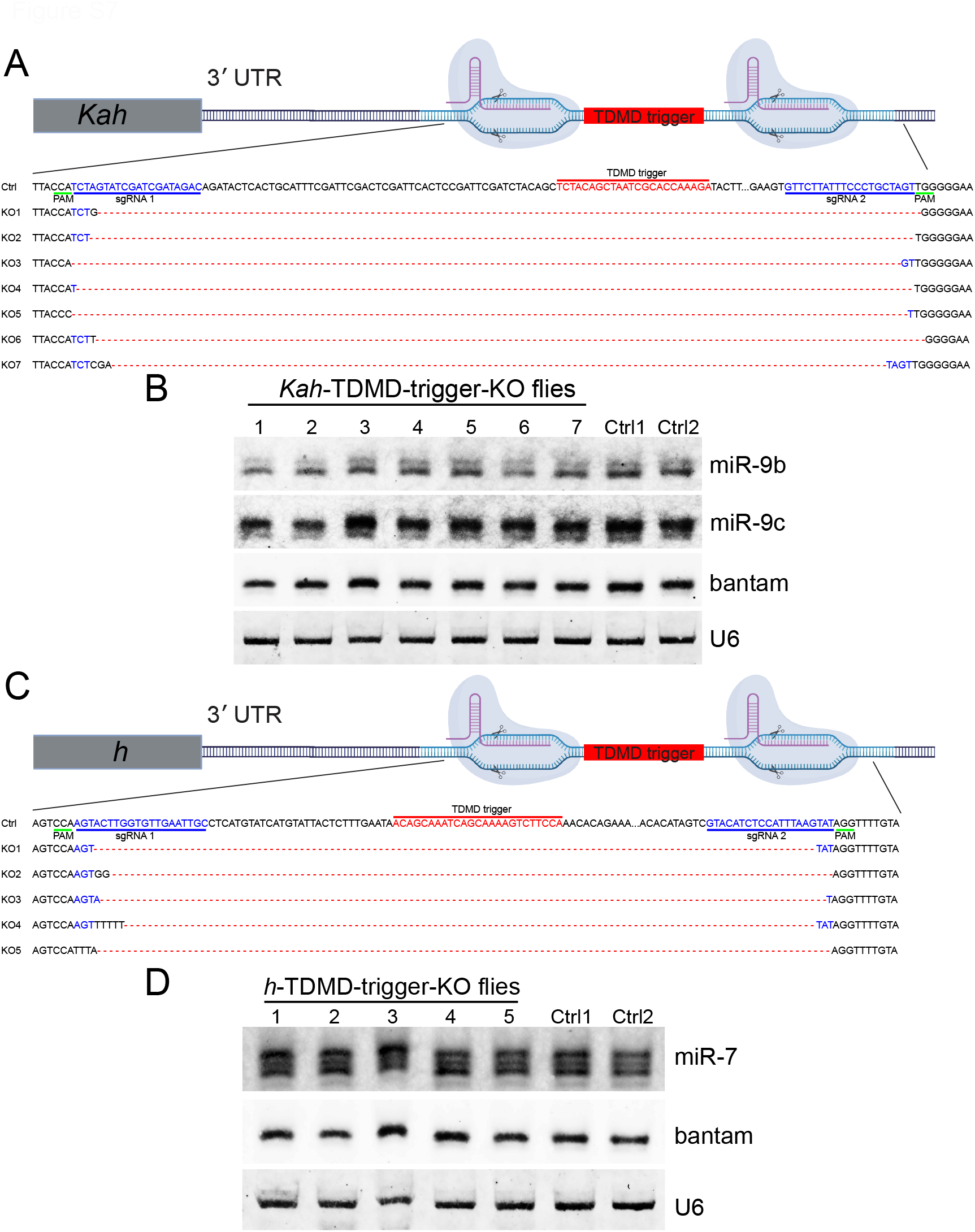
Deletion of *Kah* and *h* TDMD trigger in *Drosophila*. Schematic of the CRISPR-Cas9-mediated mutation of TDMD trigger from *Kah* 3′ UTR (**A**) and *h* 3′UTR (**C**). The TDMD trigger and sgRNAs are highlighted in red and blue, respectively. PAM sequences are underlined (green). The genotype of the control line and each mutant line is shown below. Northern blot analyses detect miR-9b (**B**), miR-7 (**D**), bantam and U6 in control lines and TDMD trigger mutant lines. Bantam and U6 serve as loading controls.

